# Azole resistance: Insights from Y132 substitutions in *Candida* sterol 14α-demethylase

**DOI:** 10.1101/2023.11.13.566813

**Authors:** R. Shyama Prasad Rao, Larina Pinto, Renuka Suravajhala, Belle Damodara Shenoy, Sudeep D. Ghate

## Abstract

**Background:** Azole-resistant *Candida* infections are on the rise. Resistant substitutions at Y132 in sterol 14α-demethylase, the key target of azole drugs, are frequent. However, it is unclear why only some Y132 substitutions are favoured or how they exert differential effects on different azoles.

**Materials and Methods:** Reported instances of Y132 substitutions were collected from the literature. Extensive molecular dynamics simulations of sterol 14α-demethylase bound to fluconazole or VT1161 (VT1) were performed, and the ligand-binding free energies were computed to quantify the effects of various Y132 substitutions on azole binding/interactions.

**Results:** Three azole-resistant substitutions, Y to C/F/H, were reported at residue position 132 in sterol 14α-demethylase. The Y132H was the most common substitution in *C. albicans*, while it was Y132F in other species. Ligand-binding free energies were -13.81 kcal/mol and -35.04 kcal/mol for fluconazole and VT1, respectively. There were differences in the ligand-binding free energies after substitutions compared to the wild type protein.

**Conclusion:** Y132F and Y132H were the most frequent substitutions in *Candida* sterol 14α-demethylase. Far higher binding free energy of fluconazole in comparison with VT1 might partly explain its susceptibility to azole-resistant substitutions. The results give key insights into azole resistance, and antifungal drug discovery and optimization.

## 1. Introduction

Fungal infections have a considerable disease burden as they affect nearly a billion people worldwide. They are estimated to be responsible for around 1.5 million deaths per year (Bongomin et al., 2017). Candidiasis, the infections caused by fungi of the *Candida* genus are becoming more prevalent. For example, vulvovaginal candidiasis affects some 138 million women annually (Denning et al., 2018). Invasive candidiasis, in particular, is becoming an emerging threat with *C. albicans* being the most common species (Pappas et al., 2018). Infections from many other *Candida* species namely *C. auris, C. glabrata, C. krusei, C. parapsilosis*, and *C. tropicalis* are also common. While azoles, the inexpensive and most frequent class of first/second-line of antifungal drugs, are effective; there are concerns of resistance to them. For example, based on a large-scale screening data, about 3.5 % of *C. albicans* isolates (n = 5265) and 96.6 % of *C. krusei* isolates (n = 1075) were found to be resistant to fluconazole (Whaley et al., 2017). Drug-resistant *Candida* infections in immunocompromised patients and the emergence of multidrug-resistant *Candida* are of particular concern (Forsberg et al., 2019).

Antifungal resistance can arise from several mechanisms. One key mechanism identified in the clinical isolates of *C. glabrata* is the disruption of a mismatch repair gene *MSH2* through mutations leading to hyper-mutable phenotype and emergence of antifungal resistant mutants (Healey et al., 2016). Other well-known mechanisms are the upregulation of the target protein production and/or efflux pumps. Thus, from the mechanistic point of view, azole resistance could be very complex (Pappas et al., 2018; Paul et al., 2022). However, as azoles target the enzyme sterol 14α-demethylase (also known as ERG11 protein, encoded by *ERG11* gene) to inhibit the biosynthesis of ergosterol and interfere with the functioning of fungal cell membrane, azole resistance is often linked to the emergence of amino acid substitutions in this target protein/enzyme.

While the minimum inhibitory concentration (MIC) breakpoint for fluconazole in susceptible *C. albicans* isolates is ≤2 μg/ml, it is ≥8 μg/ml in resistant ones (Pristov and Ghannoum, 2019). In fact, numerous resistant substitutions that strongly alter the azole MIC have been reported in sterol 14α-demethylase from the clinical isolates of many *Candida* species (Morio et al., 2010; Xiang et al., 2013). One of the most frequent resistant substitutions in sterol 14α-demethylase is at position Y132 (Flowers et al., 2015; Xiang et al., 2013). The Y132F substitution has been shown to impart a MIC of 64 to >128 μg/ml for fluconazole and 4 to >16 μg/ml for voriconazole in *C. auris* isolates (Healey et al., 2018). It is notable that many resistant substitutions are known to occur at Y132 such as Y132C, Y132F, and Y132H (Morio et al., 2010). Interestingly, different substitutions appear to have differential effects on azole resistance. For example, Y132F has a much higher MIC value (>64 μg/ml) in comparison to Y132H (16 to 32 μg/ml) for fluconazole (Xiang et al., 2013).

How substitutions, specifically Y132 substitutions, affect azole-binding is an important area of exploration. It has been suggested that substitutions lower the binding efficacy by disrupting the interactions of azoles with the target protein. For example, Y132H substitution in *C. albicans* sterol 14α-demethylase was shown to confer fluconazole resistance by preventing it from binding to haem (Kelly et al., 1999). If so, it is unclear why Y132F has a stronger effect or higher MIC value compared to Y132H (Xiang et al., 2013). Further, it is unclear why they exert differential effects on different azoles even though all azoles interact with Y132 (Hargrove et al., 2017). For example, Y132 substitutions, while effective against short-tailed azoles fluconazole and voriconazole, have no effect against medium/long-tailed azoles such as VT1 or itraconazole (Morio et al., 2010). We wanted to find whether different azoles have different binding free energies, and different substitutions differentially affect the interactions with the azoles.

We gathered the reported instances of Y132 substitutions from the literature to find their relative frequency. Further, we performed extensive molecular dynamics (MD) simulations of sterol 14α-demethylase bound to fluconazole or VT1 and computed the ligand-binding free energies using Molecular Mechanics/Poisson-Boltzmann Surface Area (MM/PBSA) method to quantify the effects of various Y132 substitutions on azole binding/interactions. The Y132H was the most common substitution in *C. albicans*, while it was Y132F in other species. We found a far higher binding free energy for fluconazole compared to VT1 that partly explain its susceptibility to azole-resistant substitutions. There were subtler differences in the azole-binding free energies upon different substitutions compared to the wild type protein. Our results are valuable in the context of azole resistance, and antifungal drug discovery and optimization.

## 2. Materials and Methods

### 2.1. Data on Y132 substitutions in sterol 14_α_-demethylase

We searched PubMed/Medline, Scopus, Web of Science, and Google Scholar databases for publications using combinations of keywords such as “azole resistance”, “*Candida*”, “mutation”, AND/OR “sterol 14α-demethylase”, and listed empirically known azole-resistant Y132 substitutions in sterol 14α-demethylase. We found 75 papers reporting 843 instances of Y132 substitutions (Table S1).

### 2.2. Acquisition of sequences and 3D structure

The sterol 14α-demethylase sequences (for *C. albicans, C. auris, C. glabrata, C. krusei, C. orthopsilosis, C. parapsilosis*, and *C. tropicalis*) were obtained from the UniProt database (https://www.uniprot.org/, last accessed on 20-10-2023). The 3D structure of *C. albicans* sterol 14α-demethylase (PDB:5TZ1) was obtained from Protein Data Bank (PDB, https://www.rcsb.org/, last accessed on 20-10-2023) in the PDB format. The PDB:5TZ1 was the highest resolution structure (2.0 Å) available for *C. albicans* sterol 14α-demethylase. The *C. albicans* sterol 14α-demethylase sequence (UniProt ID P10613) and the ligand binding residues (based on PBD:5TZ1/5ESE) are shown in Fig. 1A, and the multiple sequence alignment (based on Clustal Omega) of different sterol 14α-demethylases around Y132 ± 20 residues is shown in Fig. 1B.

**Fig. 1.**
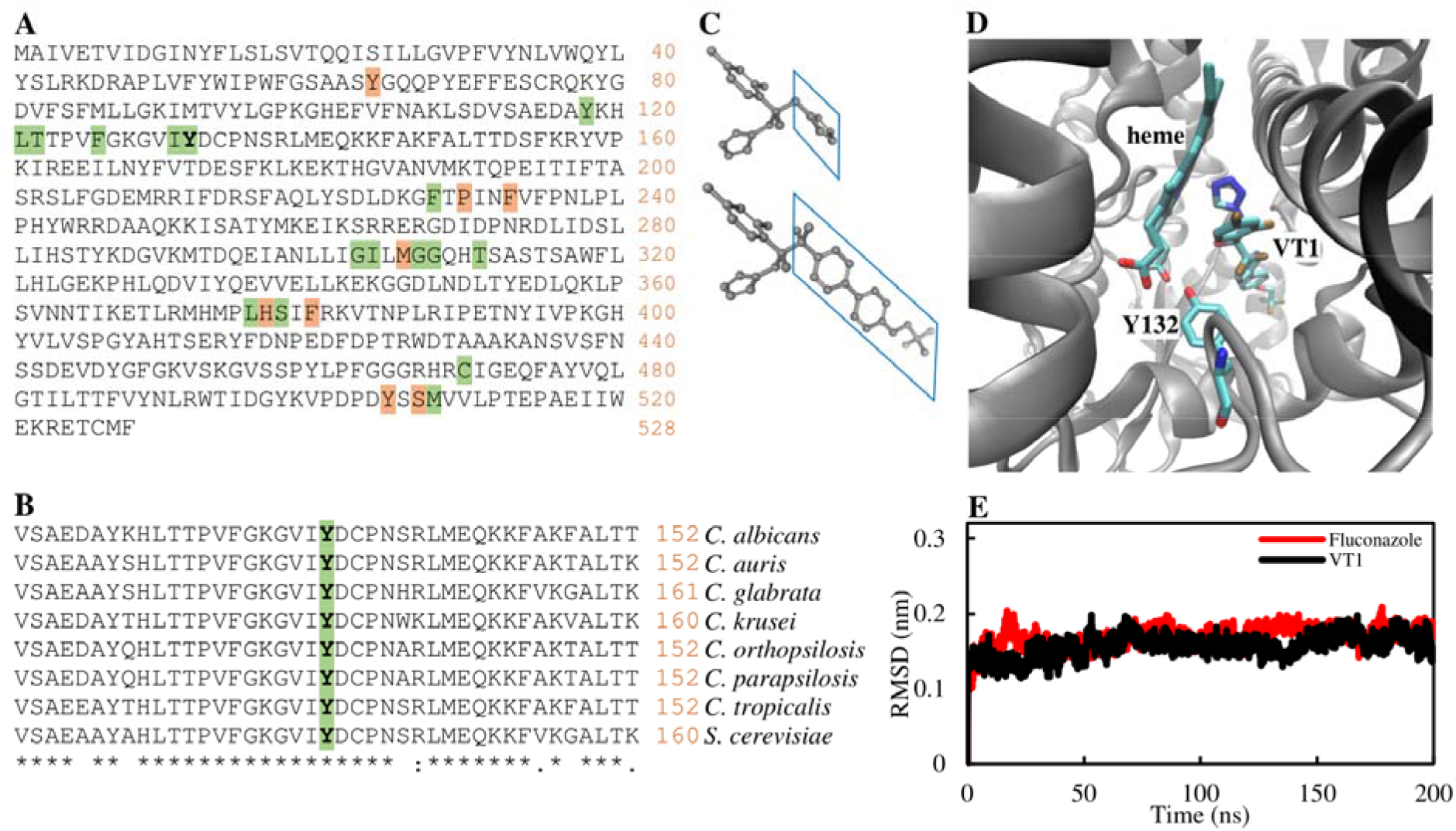
The *C. albicans* sterol 14α-demethylase. (A) The amino acid sequence (UniProt ID P10613). The residues interacting with the ligands (based on PBD:5TZ1/5ESE) are highlighted – light green for those that interact with both fluconazole and VT1, and light red for VT1 only. (B) Multiple sequence alignment (based on Clustal Omega) of sterol 14α-demethylases from different *Candida* species and *S. cerevisiae* – sequence conservation at Y132 ± 20 residues. (C) Schematic 3D structures of fluconazole and VT1 highlighting the short and medium/long-tails. (D) The 3D structural environment of Y132 (based on PBD:5TZ1) showing the proximity with heme and the ligand (VT1). (E) MD simulation – RMSD profiles of sterol 14α-demethylase complexed with fluconazole or VT1.

### 2.3. Ligand preparation

The 2D structures of two azoles – the short-tailed fluconazole and the medium/long-tailed VT1 (also known as VT1161, oteseconazole) – were downloaded from the PubChem database (https://pubchem.ncbi.nlm.nih.gov/, last accessed on 20-10-2023) in structured data file (SDF) format. The 2D structures were converted to 3D format (Fig. 1C) using Open Babel command line program (O’Boyle et al., 2011). They were finally converted to pdbqt format for docking using ‘mk_prepare_ligand.py’ script in the AutoDockFR command line software suite (Ravindranath et al., 2015).

### 2.4. Protein reparation

Chain B and water molecules present in the PDB:5TZ1 3D structure were removed using the Discovery Studio GUI software. Then the protein was protonated using the REDUCE command and finally prepared for docking using ‘prepare_receptor’ script in the AutoDockFR command line software suite (Ravindranath et al., 2015).

### 2.5. Y132 substitutions

The amino acid substitutions were modeled in the desired position (Y132) of PDB:5TZ1 using the Structure Editing/Rotamer function of ChimeraX (Goddard et al., 2018) and the best aligning rotamer was selected from the Dunbrack rotamer library (Shapovalov and Dunbrack, 2011). All atomic clashes were resolved using UCSF Chimera’s energy minimization with its default settings (steepest descent steps: 20, steepest descent step size (in angstroms): 0.02, conjugate gradient steps: 1, conjugate gradient step size (in angstroms): 0.02.). Mutated proteins were then used for molecular docking and MD simulations.

### 2.6. Molecular docking

The molecular docking of azoles to *Candida* sterol 14α-demethylase was performed in AutoDock Vina version 1.2.3 command line software (Eberhardt et al., 2021). Binding site coordinates (x, y, and z) were determined based on the cocrystal ligand VT1 in the 5TZ1 structure and prepared as a 16 Å × 16 Å × 16 Å grid by using the Receptor Grid Preparation tab in the AutoDock GUI tool. An additional grid of 40 × 40 × 40 Å was also setup for blind docking. The default values were used for other parameters while the exhaustiveness of 32 was used for better searching space (Eberhardt et al., 2021). The best-fit poses from the docking were used for MD simulations.

### 2.7. MD simulations

The protein-ligand complexes obtained from molecular docking were used to perform MD simulations using the GROMACS 2022.4 software (Van Der Spoel et al., 2005). The protein (including the heme) and the ligand parameters were generated using CHARMM36 force-field, the topology for the protein was generated using pdb2gmx utility of GROMACS, while the CGenFF online program (https://cgenff.silcsbio.com/) was used for generating the ligand topologies (Vanommeslaeghe et al., 2012). The systems (protein-ligand complexes) were prepared for MD simulations by centering them in a triclinic box with a distance of 1.0□nm from the edge and solvated with three-point transferable intermolecular potential (TIP3P) water (Bjelkmar et al., 2010). This was then followed by charge-neutralization using Cl^−^ or Na^+^ counter ions and relaxed through 50000 steps of the steepest descent algorithm for energy minimization calculations at a tolerance value of 1000□kJ/(mol.nm). The systems were equilibrated with position restraint on the protein and ligand molecules for 0.1□ns using NVT and NPT ensembles. During this process, the systems were heated to 300□K using Berendsen thermostat (Berendsen et al., 1984) with a coupling time of 2□ps and the pressure was maintained with a coupling to a reference pressure of 1□bar. Hydrogen-bonds were restrained with the LINCS algorithm (Hess et al., 1997) and the particle mesh Ewald (PME) method was used to treat the electrostatic interactions (Darden et al., 1993) for energy minimization, NVT, and NPT relaxation simulation. The cut-off radii for the van der Waals and Coulomb interactions were set at 12.0 Å. The MD simulations were performed with the integration method “Leap Frog” by using time steps of 2□fs, and trajectory snapshots were captured at every 1□ps. The MD trajectories were analysed using GROMACS utilities, and parameters such as root mean square deviation (RMSD), root mean square fluctuation (RMSF), radius of gyration (Rg), solvent-accessible surface area (SASA), and number of intermolecular hydrogen bonds (H-bonds) were chosen for analysis. The results were plotted in QtGrace tool version 0.2.6. After the stabilization of thermodynamic properties, initial MD simulations were performed for 200 ns to test the stability of the protein-ligand systems. Protein structural alignments and visualizations after docking/simulations were performed in ChimeraX (Goddard et al., 2018). The structure 2D plots to visualize the atomic interactions were done in Discovery Studio.

### 2.8. MM/PBSA calculation

The MM/PBSA method (Kumari et al., 2014; Miller et al., 2012) was used to evaluate the ligand-binding free energies and the relative stabilities of the systems. The MM/PBSA method uses the following equation:

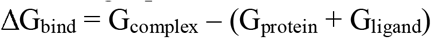

where ΔG_bind_ is the binding free energy between the protein and the ligand. It is the difference between the total free energy of the complex (G_complex_), and the sum of the free energies of the protein (G_protein_) and the ligand (G_ligand_).

The binding free energy is actually the combination of enthalpy and entropy terms as:

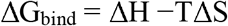

where ΔH is the enthalpy of binding, and TΔS is the entropic contribution to the free energy in a vacuum, in which T is temperature and S is entropy. Given that the entropy differences are extremely small, that term is conventionally omitted. The ΔH can be decomposed into:

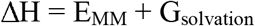

where E_MM_ is the molecular mechanics internal energy of the molecule which is the sum of electrostatic (E_ele_) and van der Waals (E_vdw_) energies. The solvation free energy (G_solvation_) is the sum of polar (G_polar_) and nonpolar (G_nonpolar_) contributions. The G_nonpolar_ is calculated from the SASA as:

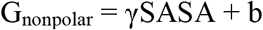

where γ = 0.0072 kcal/mol/Å, and b = 0 kcal/mol.

Specifically, the ligand-binding free energies were calculated from the GROMACS trajectory and topology files using gmxMMPBSA tool (Valdés-Tresanco et al., 2021). Due to the high computational cost, the final MD simulations for ligand binding free energy calculations were performed for 20 ns for each system (wild type or mutant protein complexed with the ligand), and a total of 100 snapshots chosen evenly from the last 10 ns (10 to 20 ns) MD trajectory was used for the calculation (Chen et al., 2019).

### 2.9. Principal Component Analysis (PCA)

PCA extracts correlated fluctuations from the MD trajectories and thus can provide information on the motion of proteins. The top few principal components (PCs) carry the most information (Maisuradze et al., 2010). The dynamics of the protein with regard to the backbone was computed using the gmx covar module of GROMACS, which builds and diagonalizes the covariance matrix to get the principal components. The first two PCs (PC1 and PC2) that explain most of the variance were plotted in Microsoft Excel.

### 2.10. Calculation of atomic distance

In order to compute the closest distance between the protein residue and the ligand, the GROMACS command “gmx distance” was used, which determines the distance between the center of mass of two atoms (Sun et al., 2020).

The average (± standard deviation) values are given for various parameters (such as binding free energy, RMSD, or distance). A t-test was performed to compared if two means (for example, the distance between sets of two atoms) are statistically different.

## 3. Results

### 3.1. Frequency of Y132 substitutions in sterol 14_α_-demethylase

Based on a through literature search, we found as many as 843 instances of azole-resistant Y132 substitutions in *Candida* sterol 14α-demethylase reported in 75 publications (Table 1 and S1). It may be noted that as these are independent publications, they provided an independent validation of the substitutions at Y132. Further, when a publication reported multiple instances of Y132 substitutions, they came from different clinical isolates/sequences. There were only three azole-resistant substitutions reported at Y132 that is Y to C, Y to H, and Y to F. Notably, Y to F was the most frequent substitution (91.8%) at Y132 in *Candida* sterol 14α-demethylase – as many as 506 instances of Y132F were found in *C. parapsilosis*, 151 instance in *C. tropicalis*, and 86 instances in *C. auris* (Table 1). However, among the 89 instances of Y132 substitutions in *C. albicans*, Y132H was the most frequent (75.3%).

**Table 1.** Y132 substitutions in sterol 14α-demethylase of different *Candida* species.

| Species | Substitution | # of instances<sup>1</sup> | # of references |
| --- | --- | --- | --- |
| *C. albicans* | Y132C | 1 | 1 |
|  | Y132F | 21 | 7 |
|  | Y132H | 67 | 21 |
| *C. auris* | Y132F | 86 | 9 |
| *C. glabrata* | Y141H<sup>2</sup> | 1 | 1 |
| *C. orthopsilosis* | Y132F | 10 | 1 |
| *C. parapsilosis* | Y132F | 506 | 29 |
| *C. tropicalis* | Y132F | 151 | 11 |
| **Total** | **-** | **843** | **75** |

<sup>1</sup>From different references or clinical isolates/sequences – see Table S1; <sup>2</sup>Y141 in *C. glabrata* corresponds to Y132 in *C. albicans*.

### 3.2. Molecular docking

The molecular docking of azoles to *Candida* sterol 14α-demethylase was done using AutoDock Vina. Even under blind docking procedure that specifies no grid, both fluconazole and VT1 docked to sterol 14α-demethylase at the lanosterol substrate-binding site that was also the known azole-binding pocket. Further, the top binding poses were similar to that of empirically known orientations (Fig. 1D, Hargrove et al., 2017). The top docking scores were -8.52 kcal/mol and -12.34 kcal/mol for fluconazole and VT1, respectively. The grid-based docking procedure yielded much lower docking scores of -9.85 kcal/mol and -15.85 kcal/mol for fluconazole and VT1, respectively. However, given that the docking scores are based on simple scoring functions (Ramírez and Caballero, 2016), they are not robust enough and thus were not used for the evaluation of binding affinities.

### 3.3. Molecular dynamics simulations

GROMACS MD simulations were performed to evaluate the stability/dynamics of the protein-ligand complexes. The RMSD of backbone or Cα atoms with respect to their initial positions indicate the stability of the system. From the initial 200 ns MD simulation trajectories (Fig. 1E), it was clear that the sterol 14α-demethylase and the ligand (fluconazole or VT1) formed stable systems. Further, the systems quickly achieved equilibrium as evident from the flat RMSD trajectories. Similarly, upon substitutions, all systems attained equilibrium within 1 to 2 ns of simulations (Fig. 2A/D and Fig. S1A/D). The Rg profile are given in Fig. 2B/E and Fig. S1B/E, and SASA profiles are shown in Fig. 2C/F and Fig. S1C/F. The wild type protein with fluconazole (but not with VT1) seemed to attain slight compaction at the end of simulation. For example, average Rg (between 10 to 20 ns) of Y132H mutant protein with fluconazole was 2.286 nm which was about 1.07% more than the wild type protein. Given the large fluctuations, there were no obvious patterns in the SASA profiles. It appears that the region at or around Y132 are less mobile as evident from the low RMSF values (Fig. S2A and S2B). The differences in the RMSF values between wild type and mutant proteins, if any, correspond to the loop regions that are bound to show random fluctuations during the simulations.

**Fig. 2.**
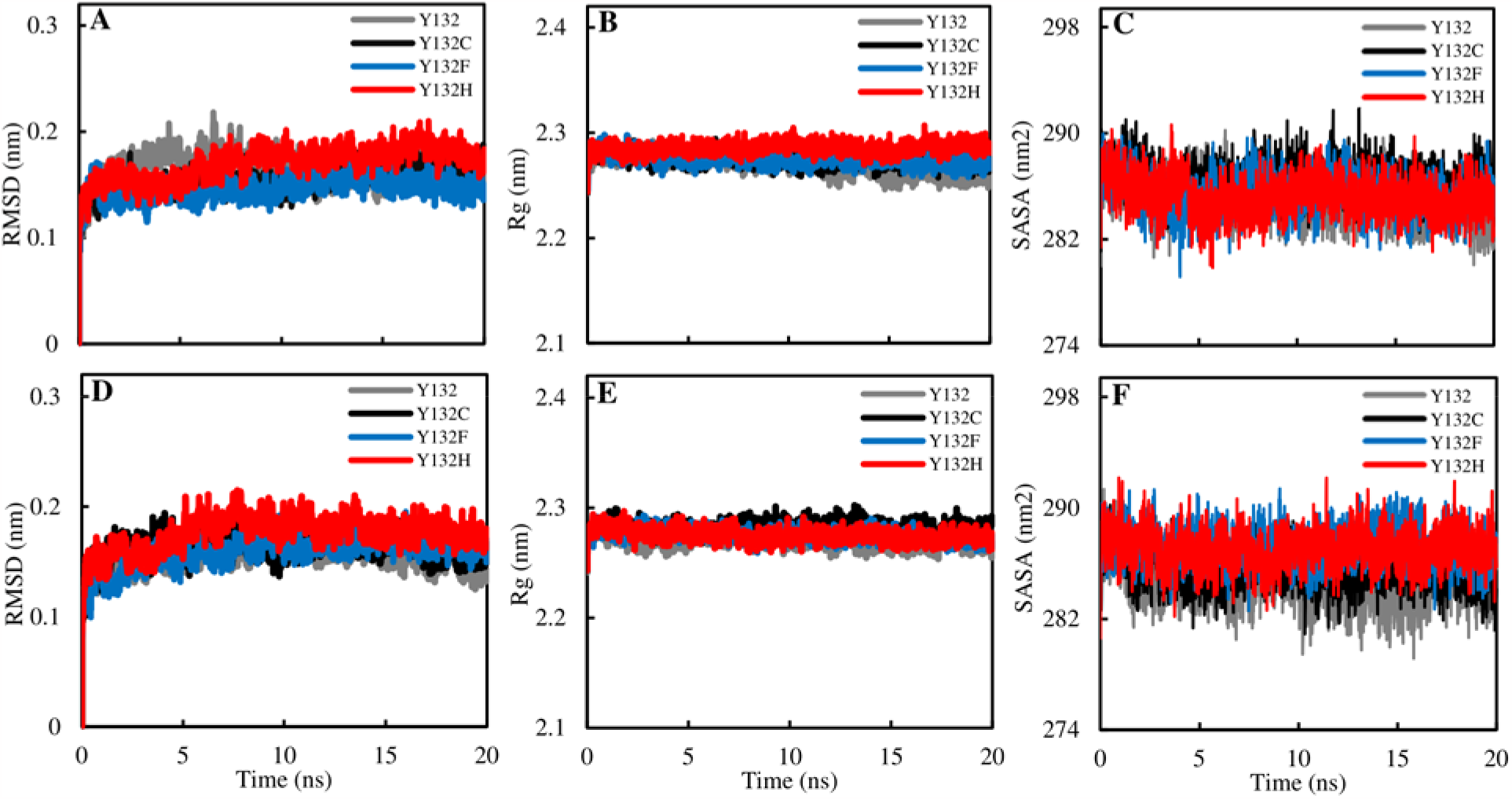
Stability analysis of ligand-bound sterol 14α-demethylase. Profiles of the backbone Cα RMSD for wild type (Y132) and mutant (Y132C, Y132F, and Y132H) proteins bound with (A) fluconazole and (D) VT1. All systems attained equilibrium within 1 to 2 ns of simulation. Profiles of the backbone Cα Rg with (B) fluconazole and (E) VT1, as well as the SASA with (C) fluconazole and (F) VT1. The wild type protein with fluconazole seemed to attain slight compaction at the end of simulation. There was no obvious difference in SASA profiles. See Fig. S1 for additional plots.

### 3.4. Ligand-binding free energies

The ligand-binding free energies of the azole-protein complexes were calculated using MM/PBSA method. The binding free energy of fluconazole with sterol 14α-demethylase wild type was -13.81 kcal/mol, while that of VT1 was -35.04 kcal/mol (Table 2). It may be noted that the ligand-binding free energy was the average of 100 snapshots chosen evenly from the last 10 ns (10 to 20 ns) MD trajectory, and therefore, has inherent fluctuations. For example, the standard deviation of binding free energy of fluconazole was ±2.19 kcal/mol and that of VT was ±2.60 kcal/mol. Further, due to the stochastic nature of MD simulations, inherent run-to-run differences in the values of free energies may be expected. The binding free energy of fluconazole with Y132H mutant protein was - 12.06, while that of VT1 was -33.53. Both in absolute and relative terms, the change in the binding free energy (between wild type versus mutant protein) was less for VT1 compared to fluconazole. The fluconazole and VT1 showed higher binding free energy with Y132C and Y132F mutant proteins (but not as much as with Y132H) compared to wild type. The substitutions that had large number of clashes or large volume such as Y132R and Y132W showed higher ligand-binding free energy compared to wild type, while others such as Y132I and Y132M showed much lower difference (Table 2).

**Table 2.** Binding free energies (ΔG) of fluconazole and VT1 with sterol 14α-demethylase under different substitutions at Y132.

| # | Amino acid | Volume ($\text{\AA}^3$)<sup>1</sup> | # of clashes<sup>2</sup> | $\Delta G$ (kcal/mol) | |
| --- | --- | --- | --- | --- | --- |
|  |  |  |  | Fluconazole | VT1 |
| **1** | **Y** | **193.6** | **-** | **-13.81** | **-35.04** |
| **2** | **C** | **108.5** | **0** | **-13.13** | **-34.83** |
| **3** | **F** | **189.9** | **2** | **-13.63** | **-34.97** |
| **4** | **H** | **153.2** | **12** | **-12.06** | **-33.53** |
| 5 | A | 88.6 | 0 | -12.33 | -35.86 |
| 6 | D | 111.1 | 1 | -13.34 | -32.99 |
| 7 | E | 138.4 | 6 | -12.97 | -32.58 |
| 8 | G | 60.1 | 0 | -13.46 | -33.72 |
| 9 | I | 166.7 | 6 | -13.96 | -33.30 |
| 10 | K | 168.6 | 8 | -11.92 | -31.27 |
| 11 | L | 166.7 | 1 | -13.63 | -32.69 |
| 12 | M | 162.9 | 4 | -14.11 | -35.04 |
| 13 | N | 114.1 | 1 | -12.12 | -33.58 |
| 14 | P | 112.7 | 2 | -11.45 | -33.84 |
| 15 | Q | 143.8 | 5 | -13.07 | -34.15 |
| 16 | R | 173.4 | 14 | -10.74 | -31.68 |
| 17 | S | 89.0 | 0 | -13.83 | -33.88 |
| 18 | T | 116.1 | 0 | -13.22 | -32.92 |
| 19 | V | 140.0 | 1 | -13.09 | -35.47 |
| 20 | W | 227.8 | 33 | -11.67 | -33.53 |

<sup>1</sup>Taken from https://www.imgt.org/; <sup>2</sup>After substitution based on ChimeraX (# of clashes were same for fluconazole and VT1); Rows (1-4) in bold are the wild type (Y) or mutants (C/F/H) currently known.

### 3.5. Structural comparison, atomic distance, and PCA

The ligands were present in the right orientation (Fig. 3A/B and Fig. S3A/B) in the binding pocket upon simulation. In comparison to residues Tyr (Y132) and Phe (Y132F), the Cys (Y132C) and His (Y132H) in particular, showed an orientation away from the ligand and the heme. This indicates that while Y132F can still be involved in the interaction with the ligand, Y132H in particular (that is the most frequent resistant-substitution in *C. albicans*) may not be able to form such interactions (Fig. 3B and Fig. S3B). This is evident from the structure 2D plots which show the residues in the proximity of ligands (Fig. 3C/D and Fig. S3C/D) wherein His (Y132H) is not seen. The distance plots between the ligand and the protein residue show clear difference (Fig. 3E and Fig. S3E).

**Fig. 3.**
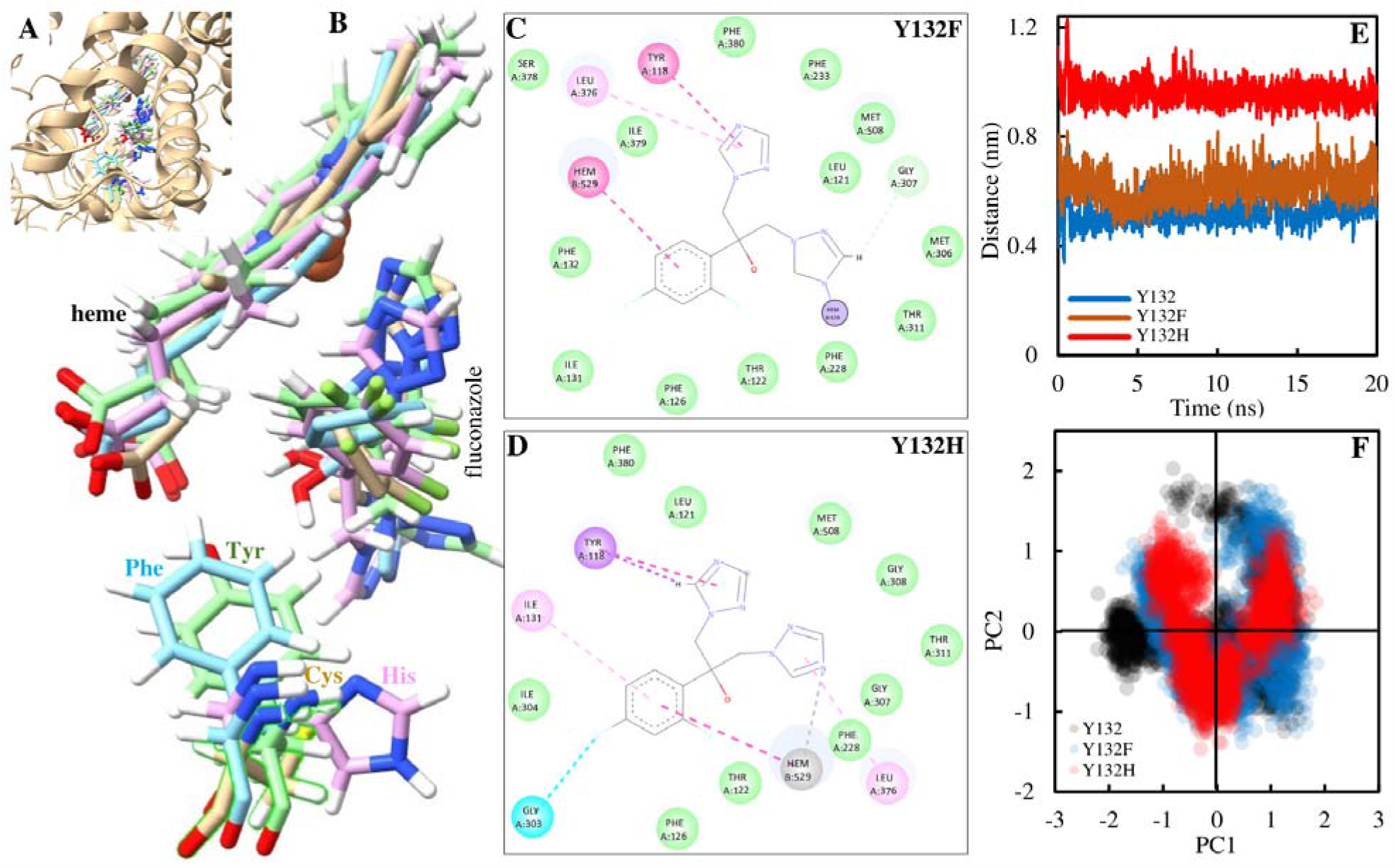
Structural alignment and comparison of substitutions at 132. (A) A close up of ligand (fluconazole) binding pocket. (B) While Phe (Y132F) has the same orientation as Tyr (Y132), Cys (Y132C) and His (Y132H) orient away from the fluconazole and the heme. The structure 2D plots confirm that while (C) Phe (Y132F) can be involved in the interaction with the fluconazole, (D) Y132H is not as it was not found in the proximity of the fluconazole. The atomic interaction colour legends are: green for van der Waals, pink for π–π stacked or alky, and blue for halogen. (E) The distance between His (Y132H) to fluconazole far greater than that of Phe (Y132F) and Tyr (Y132). (F) In comparison to wild type (Y132), the mutant proteins (Y132F and Y132H) show lower dispersion of points in the PCA plot. See Fig. S3 for additional figures/plots.

Compared to the distance between fluconazole and Tyr (Y132, 0.53±0.04 nm), the distance between fluconazole and Phe (Y132F, 0.64±0.05) was intermediate, and was very high for His (Y132H, 0.95±0.03). Similar pattern of distance was observed for VT1 with Tyr (Y132, 0.49±0.03), Phe (Y132F, 0.59±0.04), and His (Y132H, 1.02±0.05). The differences in the PCA plots (Fig. 3F and Fig. S3F) between the wild type and the mutant proteins might possibly due to overall random fluctuations in the protein rather than from specific substitutions.

## 4. Discussion

Azole-resistant *Candida* infections are increasingly becoming a public health concern (Castanheira et al., 2020). Most of the azole-resistant *Candida* isolates have been linked to substitutions at Y132 in the sterol 14α-demethylase, also known as ERG11 protein (Castanheira et al., 2020; Ceballos-Garzon et al., 2023). As evident from our compilation of data, it seems that Y132F is the only Y132 substitution in *C auris, C. parapsilosis*, and *C. tropicalis* (Castanheira et al., 2020; Ceballos-Garzon et al., 2023; Thomaz et al., 2021), whereas Y132H seems to be more frequent than Y132F in *C. albicans* (Chau et al., 2004).

While it was well known that Y132F/H substitutions confer resistance to short-tailed azoles such as fluconazole and voriconazole, but have no effect against medium/long-tailed azoles such as VT1, itraconazole, or posaconazole (Morio et al., 2010, Töpfer, 2023; Xiang et al., 2013); the reasons for such a differential action were less obvious. Sagatova et al. (2016) have shown that the Y132F/H substitutions disrupted a water-mediated hydrogen bond involved in binding of short-tailed triazoles, which contain a tertiary hydroxyl not present in long-tailed triazoles. The medium-tailed VT1 does contain that tertiary hydroxyl, yet not susceptible to Y132F/H substitution (Graham et al., 2021). From our work, it is clear that binding free energy for VT1 was far lower than that for fluconazole. Further, relative changes in the azole-binding free energies due to Y132F/H were much smaller in VT1 system compared to fluconazole. It can be argued that while any small structural perturbations due to Y132F/H substitutions might be sufficient and effective against fluconazole, they are relatively much smaller and inadequate against VT1.

It is known that Y132F and other substitutions lead to structural alteration and thus low binding efficiency between ERG11 protein and ligands (Paul et al., 2022). For example, based on molecular docking, fluconazole binding energy was shown to increase from -6.83 kcal/mol to -6.38 kcal/mol in Y132F mutant protein (Paul et al., 2022). In the current work, the Y132H substitution showed larger change in fluconazole-binding free energy compared to Y132F. This could be one of the reasons why the former is more frequent in *C. albicans* (Chau et al., 2004). On the contrary, owing to the local and global sequence differences in the sterol 14α-demethylase of other species, Y132F could be leading to a larger change in ligand-banding free energy. However, we did not test that prospect here. Another question is why only C, F, and H substitutions are preferred or observed at Y132 in sterol 14α-demethylase. Based on the free energy changes observed in this work, it is obvious that, just like Y132C/F/H, many other substitutions might have the potential to impart resistance. Those substitutions might be rarer and yet to be reported or that only Y132F/H are preferentially reported.

Another important question is whether the observed change in the binding free energy due to substitution was sufficient to impart azole resistance. There are many reports showing that numerous isolates of *C. albicans, C. auris*, and *C. parapsilosis* were indeed sensitive to azoles despite the presence of Y132F/H substitution in sterol 14α-demethylase (Castanheira et al., 2020; Chowdhary et al., 2018; Kim et al., 2022; Liu et al., 2015; Ying et al., 2013). This could also mean that Y132F/H substitution may not be effective on its own, but lends resistance in combination with other substitutions that together may cause larger structural perturbations. For example, Y132 substitution frequently occurs in combination with other substitutions such as Y132H with G450E in *C. albicans* (Chau et al., 2004; Spettel et al., 2019) and Y132F with S154F in *C. tropicalis* (Fan et al., 2023; Ngo et al., 2023). Combinations of more than two substitutions such as K128T, Y132F, and F145L, and Y132H, H283R, and G464S were also found in *C. albicans* (Chau et al., 2004). Incremental amino acid substitutions are known to impart increased inhibition (Warrilow et al., 2019). Currently we are testing the effects of multiple substitutions using computational approaches.

Our work has certain limitations. For example, we have used only *C. albicans* sterol 14α-demethylase in MD simulations. Owing to sequence/structural differences, proteins from other *Candida* species might give varying results. Further, no experimental validation being done as it is beyond the scope of this bioinformatics work.

In conclusion, Y132H was the most common Y132 substitution in sterol 14α-demethylase of *C. albicans*, while it was exclusively Y132F in many other species. The ligand-binding free energy of VT1 at -35.04 kcal/mol was far higher than that of fluconazole at -13.81 kcal/mol. There were small changes in the ligand-binding free energies after substitutions compared to the wild type protein. Far higher ligand-binding free energy and its larger relative change after substitution might partly explain the susceptibility of fluconazole compared to VT1. The results have relevance in the context of azole resistance, and antifungal drug discovery and optimization.

## Funding and Acknowledgments

This work did not receive any specific funding.

## Statement of Ethics

The work is in compliance with ethical standards. No ethical clearance was necessary.

## Data Availability

The data used in this work were obtained from literature. The relevant derived data are given in the supplemental tables.

## Author Contributions

RSPR and SDG planned and performed the work, and wrote the manuscript. LP helped in data curation. RS and BDS helped in interpretation and writing. All authors contributed intellectually, and edited/reviewed the manuscript. All authors have read and agreed to the published version of the manuscript.

## Supplemental Information

Supplemental information for this article is available online.

