## Supplemental for "Azole resistance: Insights from Y132 substitutions in *Candida* sterol 14α-demethylase"

Running title: Y132 substitutions in sterol 14α-demethylase

*Keywords: Antifungal resistance, Azole binding, Candida sterol 14α-demethylase, Molecular dynamics simulation, Sequence analysis*

**Table S1.** Y132 substitutions in sterol 14α-demethylase of different *Candida* species.

| **Species** | **Substitution** | **# of instances^1^** | **References** |
| --- | --- | --- | --- |
| *C. albicans* | Y132F | 1 | Asai et al., 1999 |
|  | Y132H | 1 | Bellamine et al., 2004 |
|  | Y132H | 13 | Chau et al., 2004 |
|  | Y132H | 2 | Debnath and Addya, 2014 |
|  | Y132H | 1 | Ding et al., 2022 |
|  | Y132H | 2 | Favre et al., 1999 |
|  | Y132F | 8 | Flowers et al., 2015 |
|  | Y132F | 1 | Goldman et al., 2004 |
|  | Y132H | 2 | Graham et al., 2021 |
|  | Y132H | 4 | Hu et al., 2015 |
|  | Y132H | 2 | Kakeya et al., 2000 |
|  | Y132H | 1 | Kelly et al., 1999 |
|  | Y132H | 1 | Kudo et al., 2005 |
|  | Y132C | 1 | Kumar et al., 2020 |
|  | Y132H | 7 | Li et al., 2004 |
|  | Y132H | 6 | Liu et al., 2015 |
|  | Y132H | 3 | Marichal et al., 1999 |
|  | Y132F/H | 3/2 | Morio et al., 2010 |
|  | Y132F | 5 | Perea et al., 2001 |
|  | Y132H | 2 | Sanglard et al., 1998 |
|  | Y132H | 2 | Spettel et al., 2019 |
|  | Y132H | 2 | Tan et al., 2014 |
|  | Y132F | 1 | Toepfer et al., 2023 |
|  | Y132H | 5 | Wang et al., 2015 |
|  | Y132F/H | 2/5 | Xiang et al., 2013 |
|  | Y132H | 2 | Xiao et al., 2004 |
|  | Y132H | 2 | Xu et al., 2008 |
| *C. auris* | Y132F | 5 | Ceballos-Garzon et al., 2023 |
|  | Y132F | 18 | Chowdhary et al., 2018 |
|  | Y132F | 24 | Healey et al., 2018 |
|  | Y132F | 21 | Hou et al., 2018 |
|  | Y132F | 1 | Lockhart, 2019 |
|  | Y132F | 6 | Rudramurthy et al., 2019 |
|  | Y132F | 1 | Rybak et al., 2021 |
|  | Y132F | 9 | Salah et al., 2021 |
|  | Y132F | 1 | Xu et al., 2023 |
| *C. glabrata* | Y141H^2^ | 1 | Vu and Moye-Rowley, 2022 |
| *C. orthopsilosis* | Y132F | 10 | Rizzato et al., 2018 |
| *C. parapsilosis* | Y132F | 87 | Alcoceba et al., 2022 |
|  | Y132F | 1 | Alobaid and Khan, 2019 |
|  | Y132F | 56 | Arastehfar et al., 2021 |
|  | Y132F | 5 | Asadzadeh et al., 2017 |
|  | Y132F | 11 | Berkow et al., 2015 |
|  | Y132F | 4 | Binder et al., 2020 |
|  | Y132F | 35 | Byun et al., 2023 |
|  | Y132F | 35 | Castanheira et al., 2020 |
|  | Y132F | 4 | Ceballos-Garzon et al., 2023 |
|  | Y132F | 9 | Choi et al., 2018 |
|  | Y132F | 7 | Corzo-Leon et al., 2021 |
|  | Y132F | 18 | Daneshnia et al., 2022 |
|  | Y132F | 33 | Demirci-Duarte et al., 2021 |
|  | Y132F | 43 | Díaz-García et al., 2022 |
|  | Y132F | 1 | Fekkar et al., 2021 |
|  | Y132F | 17 | Grossman et al., 2015 |
|  | Y132F | 1 | Hare et al., 2022 |
|  | Y132F | 1 | Hubler et al., 2023 |
|  | Y132F | 17 | Kim et al., 2022 |
|  | Y132F | 39 | Magobo et al., 2020 |
|  | Y132F | 49 | Martini et al., 2020 |
|  | Y132F | 1 | Presente et al., 2023 |
|  | Y132F | 1 | Ruma et al., 2022 |
|  | Y132F | 3 | Singh et al., 2019 |
|  | Y132F | 1 | Souza et al., 2015 |
|  | Y132F | 2 | Štefánek et al., 2023 |
|  | Y132F | 1 | Thomaz et al., 2021 |
|  | Y132F | 23 | Thomaz et al., 2022 |
|  | Y132F | 1 | Trevijano-Contador et al., 2022 |
| *C. tropicalis* | Y132F | 3 | Castanheira et al., 2020 |
|  | Y132F | 10 | Chew et al., 2016 |
|  | Y132F | 69 | Fan et al., 2023 |
|  | Y132F | 1 | Forastiero et al., 2013 |
|  | Y132F | 12 | Jiang et al., 2013 |
|  | Y132F | 11 | Jin et al., 2018 |
|  | Y132F | 21 | Ngo et al., 2023 |
|  | Y132F | 12 | Paul et al., 2022 |
|  | Y132F | 4 | Saiprom et al., 2023 |
|  | Y132F | 1 | Tan et al., 2014 |
|  | Y132F | 7 | Teo et al., 2019 |

^1^From different clinical isolates/sequences; ^2^Y141 in *C. glabrata* corresponds to Y132 in *C. albicans*.

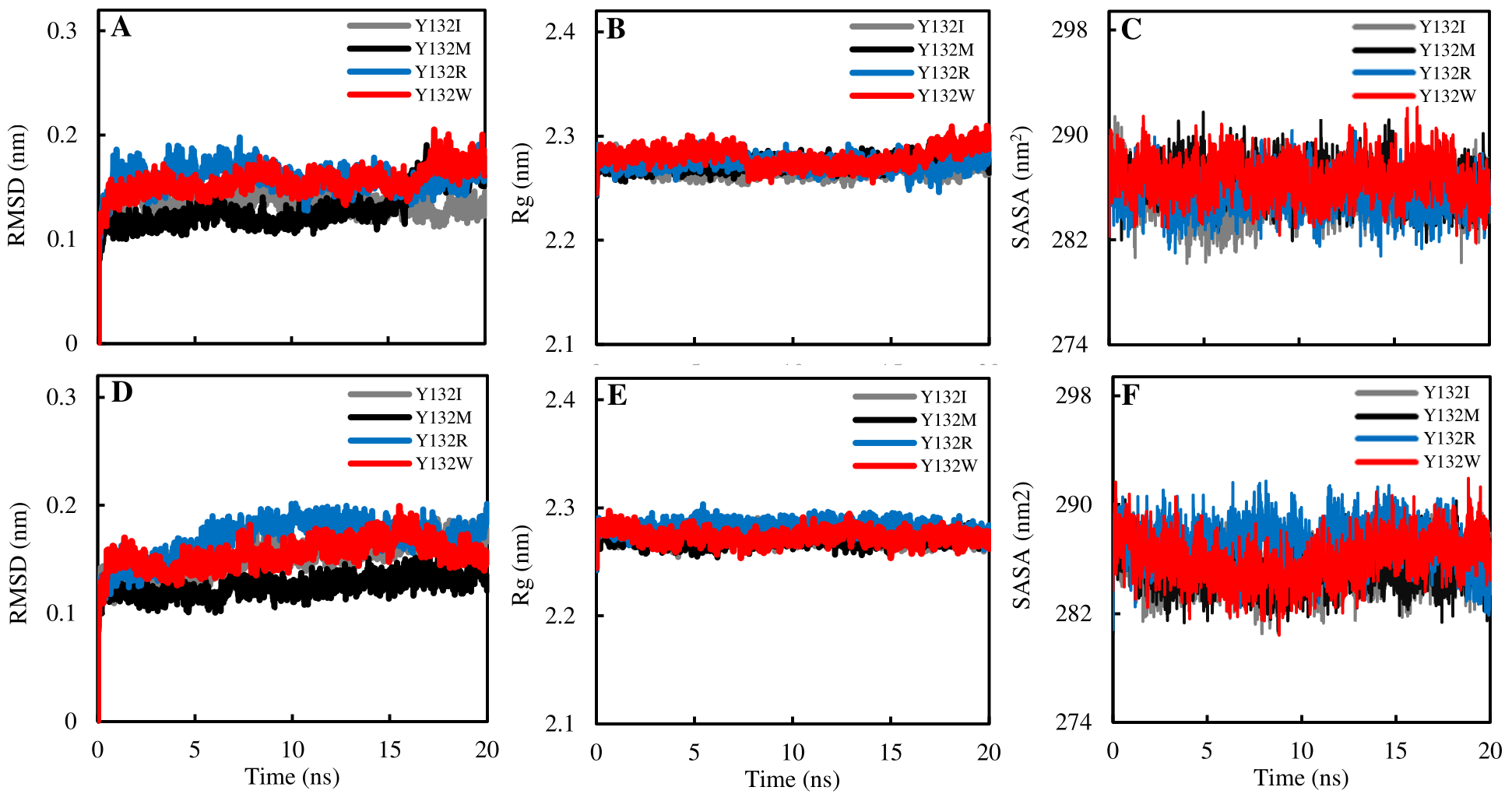

**Fig. S1.** Profiles of the backbone Cα RMSD for mutant (Y132I, Y132M, Y132R, and Y132W) proteins bound with (A) fluconazole and (D) VT1. Profiles of the backbone Cα Rg with (B) fluconazole and (E) VT1, as well as the SASA with (C) fluconazole and (F) VT1. While all systems attained equilibrium at first 1 to 2 ns of simulation, there were no obvious differences between Rg or SASA.

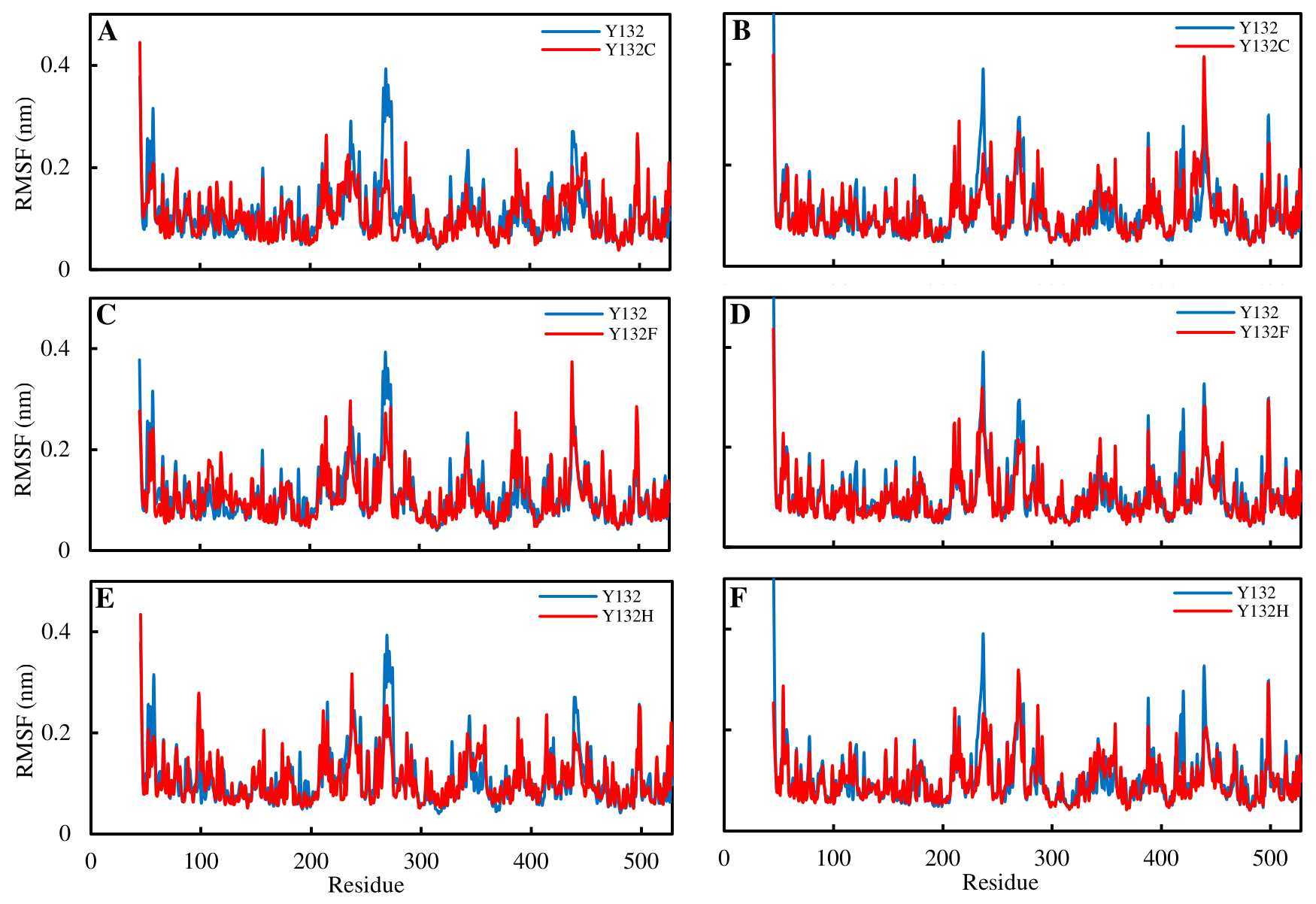

**Fig. 2A.** Comparison of RMSF values of wild type versus (A) Y132C, (C) Y132F, and (E) Y132H for fluconazole bound sterol 14α-demethylase, and wild type versus (B) Y132C, (D) Y132F, and (F) Y132H for VT1 bound sterol 14α-demethylase. The RMSF values for mutants are noticeably lower at residues 265-275 in fluconazole systems. See Fig. S2B for additional plots.

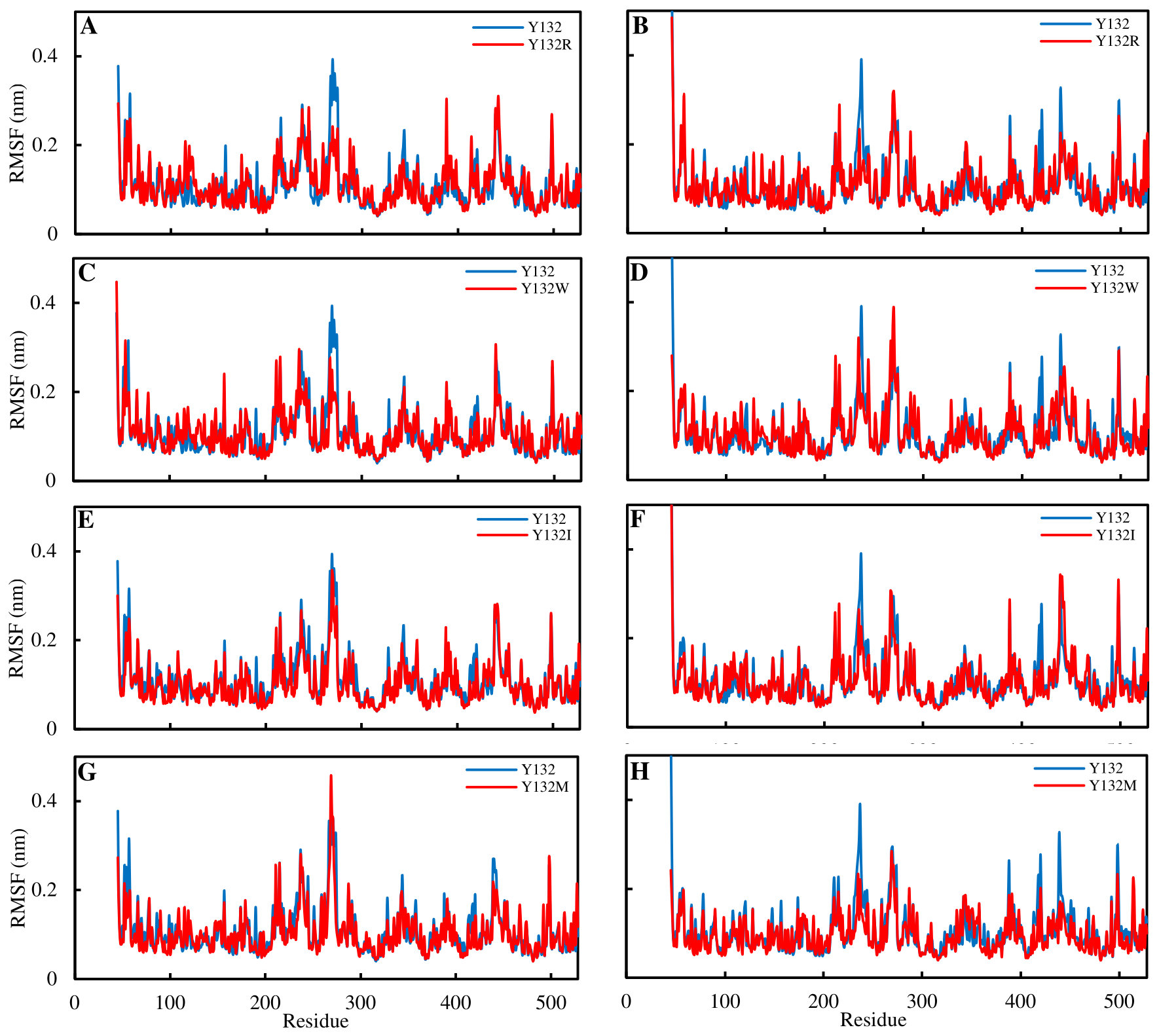

**Fig. S2B.** Comparison of RMSF values of wild type versus (A) Y132R, (C) Y132W, (E) Y132I, and (G) Y132M for fluconazole bound sterol 14α-demethylase, and wild type versus (B) Y132R, (D) Y132W, (F) Y132I, and (H) Y132M for VT1 bound sterol 14α-demethylase. The RMSF values for mutants are noticeably lower at residues 265-275 in fluconazole systems for Y132R and Y132W, but not for Y132I and Y132M. No such obvious pattern in VT1 system.

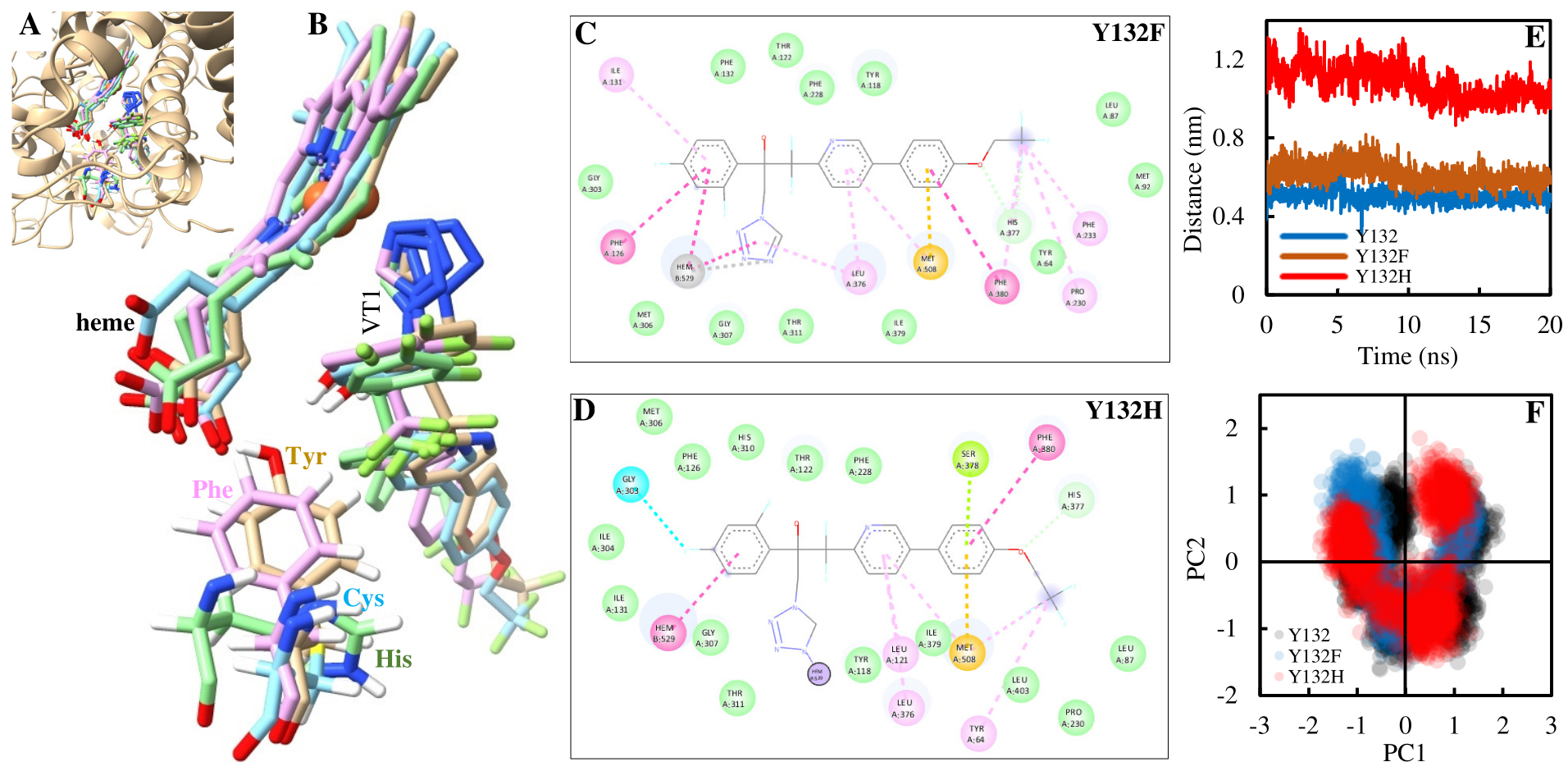

**Fig. S3.** (A) A close up of ligand (VT1) binding pocket. (B) Phe (Y132F) has the same orientation as Tyr (Y132), whereas Cys (Y132C) and His (Y132H) orient away from the VT1 and the heme. The structure 2D plots confirm that while (C) Phe (Y132F) can be involved in the interaction with the VT1, (D) Y132H is not as it was not found in the proximity of the VT1. The atomic interaction colour legends are: green for van der Waals, pink for π–π stacked or alky, and blue for halogen. (E) The distance between His (Y132H) to VT1 was far greater than that of Phe (Y132F) and Tyr (Y132). (F) In comparison to wild type (Y132), the mutant proteins (Y132F and Y132H) show lower dispersion of points in the PCA plot.
